# Integrative Species Delimitation in a Speciation Continuum: Phylogenomics, Cryptic Diversity, Diversification and Historical Biogeography of *Sinocyclocheilus* Cavefish

**DOI:** 10.1101/2025.07.09.664015

**Authors:** Yewei Liu, Tingru Mao, Jiajun Zhou, Shipeng Zhou, Yanming Peng, Hiranya Sudasinghe, Rohan Pethiyagoda, Mariana M. Vasconcellos, Marcio R. Pie, Jian Yang, Madhava Meegaskumbura

## Abstract

The transition from structured populations to distinct species often unfolds along a speciation continuum. However, empirically dissecting this continuum poses challenges for species delimitation, particularly in rapid radiations marked by recent divergence, incomplete lineage sorting, and gene flow. The species-rich *Sinocyclocheilus* cavefish radiation of Southwest China, an emerging evolutionary multi-species model system, shows a notable mismatch between its high morphological diversity and the limited divergence observed in commonly used mitochondrial DNA markers. This pattern suggests that true species richness may be underestimated. Yet, comprehensive genome-wide approaches to resolve species boundaries in this group are still lacking. To address this, we combine phylogenomics, coalescent-based species delimitation, population genetics, and historical biogeography, using both genome-wide RAD-seq and multi-locus Sanger data (including nuclear and mitochondrial DNA). Our phylogenomic analyses resolve major clades and reveal substantial cryptic diversity, well beyond what was detected by earlier markers. Species delimitation based on RAD-seq genomic data identifies several cryptic evolutionary lineages. We recover a dynamic divergence history shaped by isolation and episodic connectivity. Biogeographic reconstruction supports a mid-Miocene origin following a major vicariance event, with subsequent founder-event dispersals into subterranean habitats. This long history of fragmentation is further complicated by reticulation and ancestral polymorphism, and is reflected in present-day patterns of restricted gene flow across river valleys. These results highlight the utility of integrative genomic approaches in resolving species boundaries and uncovering the evolutionary processes that underlie high diversity in large and complex radiations.

## INTRODUCTION

Accurately delineating species, particularly within groups undergoing rapid radiation, remains challenging (De-Kayne et al., 2025). Recent and rapid divergence often leaves a complex signature of shared ancestral polymorphism and morphological convergence, which can confound taxonomic assessments, especially when based on limited data (Seehausen 2004; Nachman & Payseur 2012; Combrink et al., 2025). Distinguishing nascent species from deeply structured populations under such circumstances requires new analytical approaches beyond traditional methods. The application of genome-wide markers now provides unprecedented resolution enabling robust tests of lineage independence, and offering powerful ways of unearthing cryptic biodiversity (Edwards & Knowles 2014; Singhal et al. 2018; Streicher et al., 2024).

Cave-dwelling organisms, shaped by isolation and intense selection, provide valuable models for studying evolutionary divergence in extreme environments (Niemiller & Soares, 2014). The cyprinid genus *Sinocyclocheilus*, confined to the Karst systems of Southwest China, represents the most species-rich cavefish radiation known. With over 82 described species so far, it shows remarkable morphological disparity, including varying degrees of eye degeneration, pigmentation loss, horns on their forehead and humps on their back, and other stygomorphic features (Zhao & Zhang, 2009; Mao et al., 2021; Xiao et al., 2025; Chen et al., 2022). However, this phenotypic variation, long used for species delineation, is poorly reflected in traditional genetic markers. Many putative *Sinocyclocheilus* species exhibit low interspecific divergence in mitochondrial DNA, a discordance suggesting that past studies have likely underestimated the true species richness in this group (Mao et al. 2021, 2022).

The diversification of this genus is closely tied to the complex geological and climatic history of Southwest China (Mao et al 2025). The uplift of the Qinghai-Xizang (Tibetan) and Yunnan-Guizhou Plateau, together with cycles of aridification since the late Miocene, is thought to have fragmented subterranean habitats, promoting speciation and resulting in point endemism (Xiao et al. 2005; Mao et al., 2022). River systems have acted as both barriers and conduits in this vast karstic landscape: deep valleys often isolate basins, while underground waterways can facilitate dispersal among them, generating complex biogeographic boundaries (Wen et al., 2022). Within this geographically intricate and hydrologically complicated background, the radiation of *Sinocyclocheilus* has taken shape.

Previous taxonomic assessments on *Sinocyclocheilus* have relied primarily on mitochondrial markers and morphology (Xiao et al., 2005; Zhao & Zhang, 2009; Wen et al., 2022). These approaches have recovered relatively few genetically distinct lineages, contrasting with the striking morphological variation within the genus. Mitochondrial DNA, limited by its maternal and single-locus inheritance, often fails to capture the full complexity of divergence, particularly in systems shaped by recent radiation and incomplete lineage sorting (Hurst & Jiggins, 2005). In contrast, genome-wide approaches such as RAD-seq provide far greater resolution. When analysed within a multispecies coalescent framework, these data allow robust tests of species boundaries that incorporate gene tree discordance and ancestral polymorphism (Yang & Rannala, 2010). Yet, despite the putative high species diversity in *Sinocyclocheilus*, no comprehensive genome-wide species delimitation of the genus has yet been undertaken. As a result, its true genetically-backed species richness, including potentially extensive cryptic diversity, remains poorly characterised.

Here, we generated an extensive dataset, including mitochondrial and nuclear Sanger sequences and genome-wide RAD-seq data, building upon previous work. We aim to: (1) establish a phylogenomic framework to clarify evolutionary relationships among major clades; (2) assess species boundaries applying multiple molecular species delimitation approaches across datasets, identifying cryptic lineages and evaluating method consistency; (3) investigate population structure and gene flow to understand processes operating at the population-species interface; and (4) reconstruct the historical biogeography of the genus to elucidate the drivers of its remarkable radiation.

## 2. MATERIALS AND METHODS

We generated Sanger sequences and genome-wide RAD-seq data to investigate species boundaries and evolutionary relationships. These datasets were used to reconstruct phylogenies through multiple inference methods, followed by species delimitation analyses applied separately to each data type. Using the SNPs from the RAD-seq dataset to explore population-level processes, assess genetic clustering, and infer historical gene flow. In addition, we mapped effective migration surfaces to evaluate connectivity across the landscape. Finally, we estimated divergence times and inferred the biogeographic history using a time-calibrated phylogeny to establish the temporal and geographic context for the radiation.

### 2.1. Taxon Sampling and Molecular Data Generation

#### Sample Collection and Preservation

All experimental procedures involving live animals complied with the Chinese Animal Welfare Law (GB/T 35892-2018). Sampling of *Sinocyclocheilus* species was conducted between 2017 and 2021 across their known distribution in the karstic regions of Guangxi, Guizhou, and Yunnan provinces, China (Supplementary Fig. S1). Live fish were transported to the laboratory in oxygenated, temperature-controlled containers. Tissue samples (fin clips) were primarily collected non-lethally after anaesthetising individuals with MS-222; the specimens were allowed to recover before being returned to holding tanks. Tissue samples were immediately preserved and stored at -80°C. Our sampling aimed to include representatives from a broad range of known species and distinct morphological forms within the genus.

#### Sanger Sequencing Data

Genomic DNA was extracted from tissue samples using the DNeasy Blood and Tissue Kit (Qiagen Inc.) following the manufacturer’s protocols. Fragments of three genes, mitochondrial cytochrome b (*cytb*) and NADH dehydrogenase subunit 4 (*ND4*), and the nuclear recombination activating gene 1 (*Rag-1*), were amplified via PCR. Primers used were: DonThr R / DonGlu F for *cytb* (Sudasinghe et al., 2018), ND4F / ND4R for *ND4* (Li et al., 2008), and RAG1F1 / RAG1R1 for Rag-1 (Sudasinghe et al., 2020). PCR reactions (25 µL) contained 3 mM MgCl2, 0.4 mM dNTPs, 1X buffer, 0.06 U Taq DNA Polymerase, and 2 µM of each primer. Thermal cycling conditions included an initial denaturation at 94°C (3 min), 35 cycles of 94°C (45s), 46–50°C (1 min for *cytb*/*ND4*) or 48–56°C (1 min for Rag-1), and 72°C (45s), followed by a final extension at 72°C (5 min). PCR products were visualized on 1.5% agarose gels, purified, and sequenced in both directions using BigDye™ chemistry (Applied Biosystems Inc.) on an ABI automated sequencer. Newly generated sequences were deposited in GenBank (accession numbers provided in Supplementary Table S1). To broaden taxonomic coverage, we also incorporated 149 cytb, 126 *ND4*, and 15 Rag-1 sequences from 70 *Sinocyclocheilus* species and two outgroup taxa (*Puntius semifasciolatus* and *Cyprinus carpio*) downloaded from GenBank (Supplementary Table S1).

#### RAD-seq Data

A total of 324 individuals were sequenced using Restriction site-Associated DNA sequencing (RAD-seq), including 204 newly collected samples (representing most individuals also used for Sanger sequencing) and 120 samples previously processed by Mao et al. (2022), and 143 samples from Mao et al. (2025) (Supplementary Table S2). Two outgroup species (*Puntius semifasciolatus* and *Cyprinus carpio*) were also included. High-quality genomic DNA was extracted as described above. RAD-seq libraries were prepared following Baird et al. (2008) and related protocols. Genomic DNA was digested with the EcoRI restriction enzyme (Takara, China), adapters were ligated, and samples were pooled and sheared. Fragments ranging from 200–600 bp were size-selected from agarose gels and PCR amplified. Paired-end sequencing (100 bp) was performed on an Illumina HiSeq 2500 platform by Novogene Bioinformatics Institute (Tianjin, China).

### 2.2. Sequence Processing, Alignment, and Dataset Assembly

#### Sanger Data Processing

Forward and reverse Sanger sequences were reconciled using MAFFT v7 (https://mafft.cbrc.jp/alignment/software/mailform.html). Sequences for each gene were aligned using MAFFT v7.3.13 (Katoh & Standley, 2013) and MACSE v2.03 (Ranwez et al., 2018) as implemented in PhyloSuite v1.2.2 (Zhang et al., 2020). Alignments were manually inspected, and ambiguously aligned regions or poorly sequenced ends were trimmed using Gblocks 0.91b (Talavera & Castresana, 2007) and trimAl v1.2 (Capella-Gutierrez et al., 2009). The final concatenated dataset for phylogenetic analyses comprised 377 haplotypes from 70 *Sinocyclocheilus* species, totaling 3423 bp, partitioned by gene as follows: *cytb*: 1–1077; *ND4*: 1078–2109; *Rag-1*: 2110–3423.

#### RAD-seq Data Processing and SNP Calling

Raw RAD-seq reads were processed using ipyrad v0.9.56 (Eaton & Overcast, 2020). Reads were demultiplexed by individual, and low-quality reads (phred score <20 for >5 bases), those containing adapter contamination, or excessive poly-N tracts were discarded. A minimum sequencing depth of six reads was required for base calling. Within-individual loci were clustered using a 90% similarity threshold. To minimize paralogous loci, highly repetitive RAD tags (with stack depth exceeding a dynamically determined threshold) and their derivatives were excluded. Additionally, loci with low coverage (≤5 reads) were also removed. Several filtered SNP datasets were generated from the assembled RAD loci for different downstream analyses (Supplementary Table S3): i) For concatenated phylogenomic analyses (IQ-TREE, MrBayes), we used loci present in at least 90% of all 324 individuals (including outgroups); ii) For SVDquartets (tetrad), one random SNP per locus present in all individuals was selected to minimize linkage; iii) For BPP species delimitation analyses, outgroups were excluded. Because of high computational demands, two datasets were assembled: initial six clade-specific datasets (Clades A–F, see results), each retaining loci shared across all individuals within each clade, and a combined dataset of all 318 ingroup individuals, including loci present in ≥90% of all individuals. This two-step approach streamlined the final analysis by constraining parameter space within clades based on initial results. iv) For population structure (STRUCTURE, DAPC), admixture (TreeMix) and Flexible Estimation of Effective Migration Surfaces (FEEMS) analyses, outgroups were excluded, and one SNP per locus present in at least 90% of the 318 ingroup individuals were used; v) For SNAPP divergence time estimation, loci shared among all 56 species (represented by selected individuals) were used, from which unlinked SNPs were selected.

### 2.3. Phylogenetic Inference

#### Sanger Data Phylogenetic Analyses

Optimal substitution models and partitioning schemes for the concatenated Sanger datasets (*cytb*+*ND4*+*Rag-1*; and *cytb*+*ND4*) were determined using ModelFinder (Kalyaanamoorthy et al., 2017) in PhyloSuite, based on the Bayesian Information Criterion (BIC), applying a codon-partitioned model. Maximum Likelihood (ML) phylogenies were inferred using IQ-TREE v1.6.8 (Nguyen et al., 2015) with 1000 ultrafast bootstrap replicates. Bayesian Inference (BI) was performed using MrBayes v3.1.2 (Ronquist et al., 2012), running two independent analyses for 200 million generations, sampling every 20,000 generations, discarding the first 10% as burn-in. Convergence was assessed using Tracer v1.7.1 (Rambaut et al., 2018). Individual gene trees were also estimated.

#### RAD-seq Phylogenomic Analyses

Several phylogenomic approaches, including concatenated and coalescent, were applied to the RAD-seq SNP datasets: i) A ML tree was inferred from the concatenated RAD-seq loci (Dataset i) using IQ-TREE v2.2.6 (Minh et al., 2020). Support was assessed with 1000 ultrafast bootstrap replicates. ii) Bayesian inference was conducted on the same concatenated RAD-seq loci (Dataset i, excluding invariant sites) using MrBayes v3.1.2. Two independent runs of 48 MCMC chains were performed for 20 million generations, sampling every 1000 generations, with convergence diagnostics assessed as above. ModelFinder identified best-fit nucleotide substitution models for both analyses (ML and BI). iii) To account for incomplete lineage sorting (ILS) using a summary method, a species tree was inferred with SVDquartets (Chifman & Kubatko, 2015) implemented in tetrad v.0.7.19 within ipyrad API, based on the unlinked SNP dataset (Dataset ii). All possible quartets were evaluated, with support assessed from 1000 non-parametric bootstrap replicates. iv) ASTRAL-III v5.7.3 (Zhang et al., 2018) was used to infer a species tree from individual gene trees generated for each RAD locus (treated as separate genes and inferred using IQ-TREE). ASTRAL accounts for ILS and provides local posterior probabilities for branch support.

### 2.4. Species Delimitation Analyses

To confidently assess species boundaries and identify potential cryptic lineages, multiple species delimitation methods were applied to the Sanger and RAD-seq datasets to assess species boundaries and identify potential cryptic lineages.

#### Sanger Data-Based Species Delimitation

A suite of methods was employed to evaluate lineage boundaries based on traditional mitochondrial and nuclear markers: i) Automatic Barcode Gap Discovery (ABGD) was applied to *cytb*, *ND4*, and concatenated datasets via the ABGD web server (Puillandre et al., 2012), with prior intraspecific divergence (P) from 0.001 to 0.1, 10 steps, minimum relative gap width (X) of 1, and the Kimura (K80) model. ii) Bayesian Poisson Tree Processes (bPTP) was performed on the bPTP web server (Zhang et al., 2013) using ML trees (*cytb*, *ND4*, concatenated) as input. Analyses ran for 5 million MCMC generations with thinning of 100, 0.1 burn-in, and outgroups removed.iii) Multi-rate Poisson Tree Processes (mPTP) was applied to the same ML input trees on the mPTP web server (Kapli et al., 2017), with 50 million MCMC generations, sampling at every 1 million, and a burn-in of 1 million.iv) Bayesian General Mixed Yule-Coalescent (bGMYC) was implemented in R (Reid & Carstens, 2012) using 1000 post-burn-in ultrametric trees from the BEAST analysis of the Sanger data (section 2.6). Delimitation was assessed across several conspecificity probability thresholds (0.95–0.99).

#### RAD-seq Data-Based Species Delimitation (BPP)

Species delimitation using SNP data was performed with Bayesian Phylogenetics and Phylogeography (BPP) v4.3 (Yang & Rannala, 2010, 2014), which implements the multispecies coalescent model to assess species boundaries considering ILS.Analyses were conducted on Dataset iii. Due to computational constraints, initial runs focused on delimiting species within the six major clades identified through phylogenomic analyses, followed by a comprehensive analysis of all 318 *Sinocyclocheilus* individuals based on those initial results. A guide tree derived from the SVDquartets analysis was used. Priors for population size (θ) and root divergence time (τ0) were set based on Minimalist BPP (https://brannala.github.io/bpps/#/), using IG (3, 0.0025) for θ and IG (3, 0.20) for τ0, following Yang (2015). Both rjMCMC algorithm 0 (ɛ=2) and algorithm 1 (α=2, m=1) were tested across four independent runs of 20,000 MCMC steps each (sampling every 2 steps, with 2000 steps burn-in). Populations were designated as cryptic species if delimited through the RAD-seq-based BPP analysis and showed subtle morphological differences from their confirmed sister species (to be further examined in formal species descriptions).

### 2.5. Population Genetic Structure and Historical Gene Flow Analyses (using RAD-seq data)

To investigate intraspecific genetic structure, interspecific admixture, and historical gene flow, we used RAD-seq SNP data (Dataset iv) in three complementary approaches. First, we used STRUCTURE v2.3.4 (Pritchard et al., 2000) with an admixture model with correlated allele frequencies, testing K values from 2 to 57 (10 replicates per K), with 20,000 generations following a 10,000 generation burn-in. Results were summarised with STRUCTURE HARVESTER (Earl, 2012), CLUMPP (Jakobsson & Rosenberg, 2007) and distruct v1.1 (Rosenberg, 2004). Second, a Discriminant Analysis of Principal Components (DAPC) was performed using the R package adegenet v2.1.5 (Jombart, 2008), with major clades identified from phylogenomic analyses used as priori groupings. Third, we used TreeMix v1.1.3 (Pickrell & Pritchard, 2012) to infer population splits and patterns of historical gene flow based on allele frequency data, as implemented in ipyrad API. We tested the number of migration edges (m) from 1 to 25, and selected the best-fitting model based on likelihood scores.

### 2.6. Divergence Time Estimation

Divergence times based on Sanger data (*cytb*+*ND4*+*Rag-1*; and *cytb*+*ND4*) were estimated using BEAST v2.7.7 (Suchard et al., 2018). A partitioned codon model was used with substitution models selected by ModelFinder. Secondary calibrations from Li et al. (2021) included divergence between: *S. maitianheensis* and *S. anophthalmus* (mean = 2.7 Mya, SD = 1.05, 95% HPD = 1.4 to 4.8 Mya), between *S. maitianheensis* + *S. anophthalmus* and *S. grahami* (mean = 4.3 Mya, SD = 1.30, 95% HPD = 2.5 to 6.9 Mya), among *S. maitianheensis* + *S. anophthalmus* + *S. grahami* and *S. anshuiensis* (mean = 7.8 Mya, SD = 1.95, 95% HPD = 5.0 to 11.7 Mya), and among *S. maitianheensis* + *S. anophthalmus* + *S. grahami* + *S. anshuiensis* and *S. rhinocerous* (mean = 8.9 Mya, SD = 2.1, 95% HPD = 5.8 to 13.1 Mya). A Yule speciation process with constant population size and an uncorrelated lognormal molecular clock was assumed. Analyses were run for 200 million generations (sampling every 20,000), with convergence checked in Tracer v1.7.1.

To estimate divergence times from genomic data while accounting for ILS, SNAPP v1.4.1 (Bryant et al., 2012) implemented in BEAST v2.6.6 was used. Analyses were conducted on Dataset v (774 unlinked SNPs across 56 individuals representing major lineages). The input XML file was prepared using snapp_prep.rb (Stange et al., 2018). The calibration points described above were applied as priors on the corresponding node. SNAPP analyses ran for 3 million MCMC generations (sampling every 300 steps), with the first 10% discarded as burn-in. Convergence was assessed in Tracer v1.7.1, and the MCC tree was generated using TreeAnnotator v2.7.3.

### 2.7. Biogeography

#### Biogeographic Reconstructions

To investigate the geographic patterns of speciation and dispersal among *Sinocyclocheilus* species, we conducted biogeographic reconstructions in BioGeoBEARS (Matzke, 2014) implemented within the RASP (Reconstruct Ancestral State in Phylogenies) framework (Yu et al., 2020). The Karst habitats occupied by *Sinocyclocheilus* cavefish are characterised by significant isolation, which inherently limits dispersal between major hydrological basins (Ma et al., 2023). Additional factors such as the size of core distribution areas, regional geological activity, and the paleo-drainage history of river networks, also known to influence species distributions (Lei, 2023), guided the definition of nine discrete operational biogeographic areas, corresponding to major river systems currently occupied by *Sinocyclocheilus*: Guijiang/Hejiang (GH), Liujiang (LJ), Hongshuihe (HS), Nanpanjiang (NP), Beipanjiang (BP), Red River (RR), Jinsha River (JS), Wujiang (WJ), and Zuojiang (ZJ). Given the narrow distribution ranges and high endemism of most species, suggesting limited dispersal across major basins, we constrained the ancestral and extant range sizes to a maximum of two areas.

Three statistical models were explored: Dispersal-Extinction-Cladogenesis (DEC) (Ree & Smith, 2008), Dispersal-Vicariance Analysis-like (DIVALIKE) (Ronquist, 1997), and BayArea-like (BAYAREALIKE) (Landis et al., 2013). Each model was tested with and without the founder-event speciation parameter (+J) (Matzke, 2013, 2014). The "+J" parameter explicitly accounts for founder-event or "jump dispersal" speciation, a mechanism plausible in fragmented environments like Karst systems, where small, isolated populations may rapidly diverge following colonization of new cave habitats. Incorporating this parameter is therefore critical for accurately reconstructing the biogeographic history of *Sinocyclocheilus*. The time-calibrated phylogenetic tree produced in SNAPP was used as the input topology for these analyses in RASP. All the six models were compared to identify the most suitable model for *Sinocyclocheilus* and to evaluate whether incorporating the +J parameter significantly improved model fit relative to the corresponding null models (DEC, DIVALIKE, BAYAREALIKE). Model selection was based on the corrected Akaike Information Criterion (AICc) (Anderson & Burnham, 2004).

#### Estimation of Genetic Connectivity and Barriers to Gene Flow

To identify connectivity corridors and potential barriers to gene flow at a fine spatial scale, we applied the Fast and Flexible Estimation of Effective Migration Surfaces (FEEMS) method (Marcus et al., 2021). The input dataset consisted of SNPs, filtered to exclude variants with a minor allele frequency (MAF) below 5% or missing data exceeding 10%. Filtering was performed using PLINK v.1.9.0 (Chang et al., 2015), resulting in a final dataset of 23,184 SNPs across 318 individuals. Sampled populations were assigned to vertices of a customized 5 km triangular discrete global grid (DGG), generated with the R package ‘dggridR’ (Barnes et al., 2017). Effective migration rates across this grid were estimated using a penalized likelihood framework that incorporates a smoothing parameter (λ) to control model complexity. Because the estimated migration surface is sensitive to the choice of λ, where low λ values may cause overfitting and spurious inferences (Marcus et al., 2021), we employed the ‘leave-one-out’ cross-validation procedure implemented in FEEMS to identify an optimal value. This procedure tested 20 values evenly spaced across a range from 10^−6^ to 10^2^. The model corresponding to the λ value that minimized the L2 error was selected for visualization and interpretation of effective migration surfaces.

## 3. RESULTS

Our integrative analyses, combining Sanger sequencing and RAD-seq phylogenomics, for a comprehensive inference of species delimitation, population genetics and biogeography, revealed a complex evolutionary history for *Sinocyclocheilus*, characterized by substantial cryptic diversity and varying degrees of congruence among molecular datasets and analyses.

### 3.1. Sanger Data Analyses

#### Phylogenetic Relationships from Sanger Data

Phylogenetic analyses (ML and BI) of the concatenated three-gene Sanger dataset (*cytb*+*ND4*+*Rag-1*; 377 haplotypes, 3423 bp) and a two-gene mitochondrial dataset (*cytb*+*ND4*) generally provided strong support for the major clades within *Sinocyclocheilus* (Fig. 1, Supplementary Figs. S2-S5). Across both datasets, five major clades (Clades I-V) were recovered. The *S. jinxiensis* was a sister species of *S. tianlinensis* and *S. anatirostris* in ML and MrBayes trees. While the topologies of the ML and Bayesian trees were largely congruent in recovering these major clades, some topological inconsistencies were observed. For instance, the placement of species such as *S. ronganensis*, *S. macrolepis, S. angularis, S. zhenfengensis*, and *S.* cf*. furcodorsalis* differed between ML and Bayesian trees, and between the mitochondrial and the ‘mitochondrial + nuclear’ datasets (Fig. 1, Supplementary Figs. S2-S5). Relationships among certain recognised species, such as populations attributed to *S. cyphotergous* and *S. multipunctatus*, were unstable or non-monophyletic. Notably, the *Rag-1* nuclear gene tree exhibited marked topological differences compared to the mitochondrial gene trees (Supplementary Figs. S6-S7), suggesting potential discordance due to processes like incomplete lineage sorting or mitochondrial introgression.

**Figure 1.**
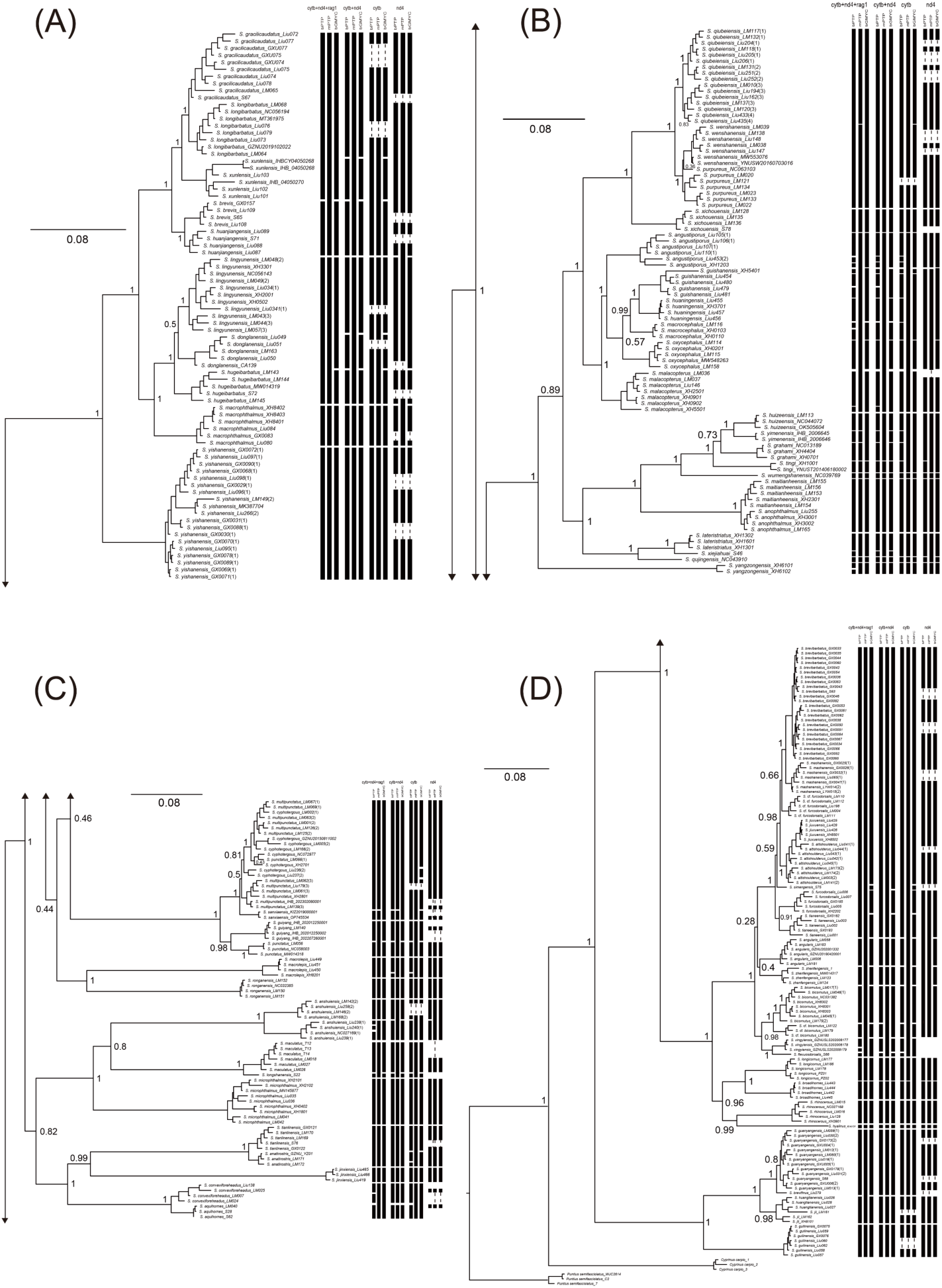
**A-D**) Molecular phylogenetic relationships of *Sinocyclocheilus* based on Bayesian inference of the concatenated cytb +nd4+ rag1 dataset. Numbers at the nodes indicate Bayesian posterior probabilities. Scale bar represents number of substitutions per site. Species delimitation results from molecular methods (bPTP, mPTP, and bGMYC) applied to cytb, ND4, Rag-1, and the concatenated dataset are shown as black rectangles to the right of the trees.

#### Species Delimitation with Sanger Data

Species delimitation methods applied to the Sanger datasets (ABGD, bPTP, mPTP, bGMYC) consistently recovered fewer putative species than the number of morphologically recognized species included in the analyses (Fig. 1, Supplementary Table S4). For the concatenated *cytb*+*ND4*+*Rag-1* dataset, these methods identified between 29 (ABGD) and 63 (bPTP) putative species. Similar ranges were observed for the *cytb*+*ND4* dataset (26-61 species) and for the individual mitochondrial markers (*cytb*: 29-55; *ND4*: 27-37 species). Most importantly, these Sanger-based delimitation approaches generally failed to consistently resolve closely related, morphologically distinct species. In several cases, they also failed to detect distinct morphological lineages that were later recovered using RAD-seq data.

### 3.2. RAD-seq Data Analyses

#### Phylogenomic Reconstruction from RAD-seq Data

RAD sequencing yielded an average of ∼3.6 billion raw reads per sample (error rate ∼0.03%; Supplementary Table S5). Phylogenomic analyses of the filtered RAD-seq SNP dataset (87,141 SNPs from loci present in ≥90% of 324 individuals) using both concatenation (IQ-TREE, MrBayes) and coalescent-based methods (SVDquartets/tetrad, ASTRAL) produced largely congruent and well-resolved topologies. These analyses consistently identified six well-supported major clades (Clades A-F; Fig. 2, Supplementary Figs. S8-S10), with most nodes receiving strong support (bootstrap values for IQ-TREE >95 and posterior probabilities >0.9 for MrBayes, ASTRAL). The ASTRAL species tree, which accounts for incomplete lineage sorting, showed maximal support (posterior probability of 1.0) for nearly all nodes (Supplementary Fig. S9). We recognized four cryptic species (*S.* cf*. bicornutus*, *S.* cf*. furcodorsalis*, *S.* cf*. qiubeiensis*, and *S.* cf*. yishanensis*), which along with their respective sister species, were strongly supported across all four phylogenetic analyses (IQ-TREE, MrBayes, ASTRAL, and tetrad). Two additional cryptic species, *S.* cf. *guanyangensis* and *S.* cf. *multipunctatus*, were strongly supported as distinct from their respective sister taxa by IQ-TREE and MrBayes analyses, however, reciprocal monophyly was not achieved in the MrBayes and Tetrad trees. Minor topological incongruences were observed between the concatenation-based (IQ-TREE; Fig. 2) and coalescent-based (ASTRAL; Supplementary Fig. S9) approaches, for example, in the placement of *S. jii*, *S. huangtianensis*, and *S. brevifinus*. The SVDquartets analysis also placed some individuals outside of their conspecific groupings, including *S. lingyunensis* (Liu172) and certain *S.* cf. *multipunctatus* and *S. guanyangensis* individuals (Supplementary Fig. S10), suggesting complex population histories or possible misidentifications. The *S. jinxiensis* is classified into Clade B with *S. aquihornes*, *S. tianlinensis* and *S. anatirostris* (IQ-TREE, MrBayes, ASTRAL, and tetrad).

**Figure 2.**
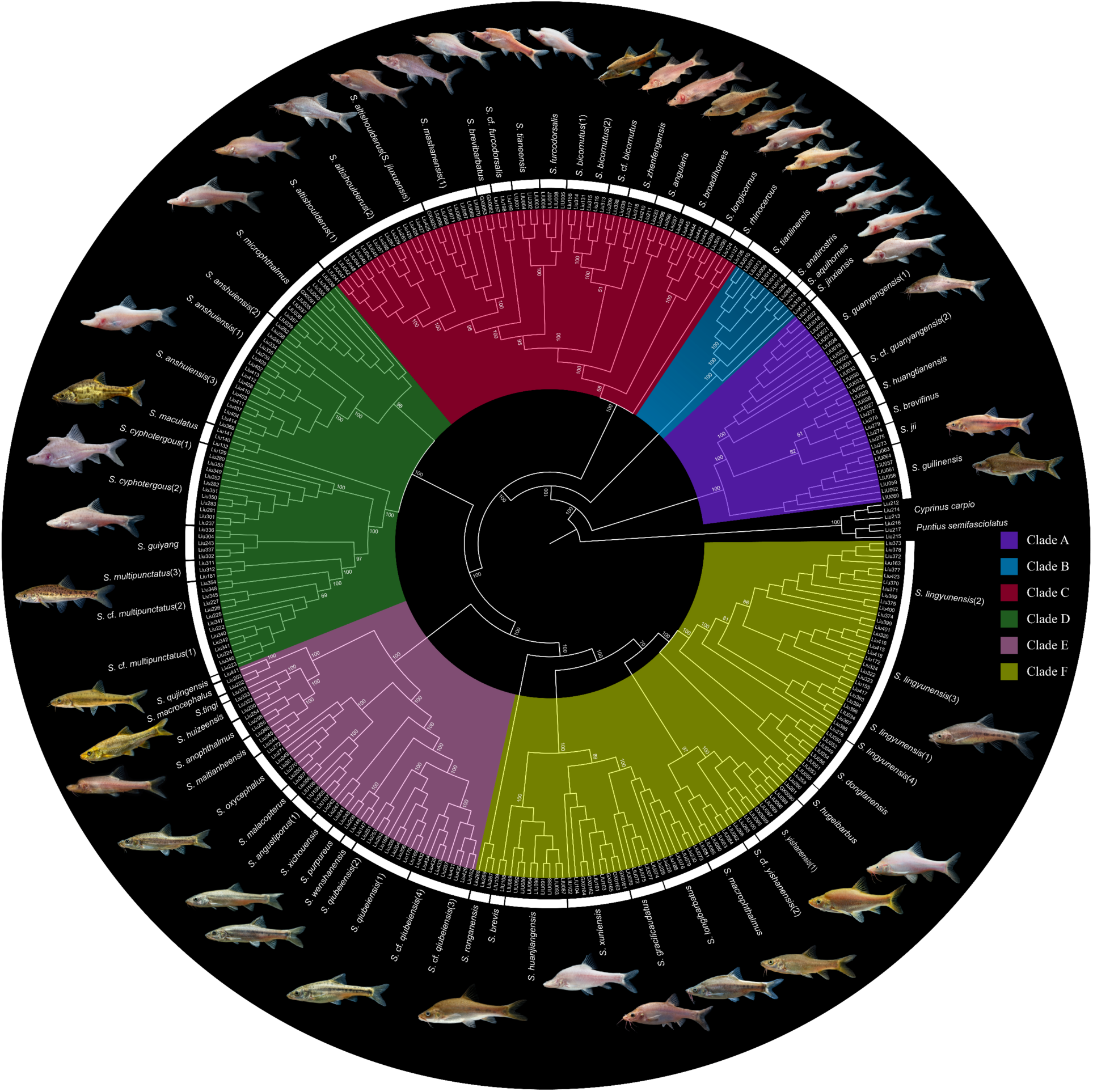
Maximum-likelihood (IQ-TREE) phylogeny of *Sinocyclocheilus* based on 87,141 RAD loci. Bootstrap support values are shown at each node. Numbers in parentheses after species names denote individuals from distinct caves (i.e. separate populations). Species delimitation results from BPP using RAD loci are depicted as white rectangles adjacent to the phylogeny.

#### Species Delimitation with RAD-seq Data (BPP)

Bayesian species delimitation using BPP on the RAD-seq SNP dataset provided strong support for a greater number of distinct evolutionary lineages compared to the Sanger dataset. The results largely agreed with the major clades identified in the phylogenomic analyses, while also further subdividing them (Supplementary Table S6, Supplementary Fig. 2). Overall, BPP analyses on the 318 ingroup *Sinocyclocheilus* individuals supported the existence of at least 56 distinct evolutionary lineages, many of which correspond to already recognized species, but several of which represent strong candidates for undescribed taxa. Within Clade A, six species were confirmed (PP = 1.0), and a population previously identified as *S. guanyangensis* were consistently delimited as a distinct potential cryptic species (*S.* cf. *guanyangensis*). Clade B comprised four species (PP=1.0). Clade C resolved into thirteen species (PP=0.8), with samples of *S. jiuxuensis* and *S. altishoulderus* forming distinct population-level clusters within a broader *S. altishoulderus* lineage. This clade also included two strongly supported candidate new species: *S.* cf. *bicornutus* and *S.* cf. *furcodorsalis*. In Clade D, seven species were delimited with strong support (PP=1.0), including two populations previously identified as *S. multipunctatus*, which were consistently recognized as a distinct putative species (*S.* cf. *multipunctatus*). Clade E resolved into fourteen species (PP=1.0), including a distinct population within *S. qiubeiensis* (S. cf. “qiubeiensis”). Within Clade F, twelve species were delimited (PP=1.0), with structured populations of *S. lingyunensis*, and one population of *S. yishanensis* forming another candidate species *(S.* cf. yishanensis.

### 3.3. Population Genetics and Evolutionary Dynamics

#### Divergence Time Estimates

Time-calibrated phylogenetic analysis using SNAPP on the RAD-seq SNP dataset estimated the crown age of the sampled *Sinocyclocheilus* to be approximately 17.96 Ma (95% HPD: 11.87-24.8 Ma) (Supplementary Fig. S11). Other recent divergence events included the subclade containing *S. brevibarbatus*, *S. mashanensis*, and *S.* cf*. furcodorsalis*, with a tMRCA of 0.8 Ma (95% HPD: 0.41-1.34 Ma). The overall topology of the SNAPP tree was congruent with the six major clades recovered in the other RAD-seq analyses.

#### Genetic Structure

STRUCTURE analysis of the RAD-seq SNP dataset for all ingroup individuals identified an optimal K=6, based on the Evanno method (Evanno et al., 2005) (Supplementary Table S7). These six genetic clusters corresponded closely to the six major phylogenomic clades (Clades A-F) recovered in previous analyses (Fig. 3B). DAPC analysis, using these six clades as a priori groups, also revealed clear differentiation among them (Fig. 3C). Clades B and D emerged as the most genetically distinct, while Clades A, C, E, and F showed comparatively lower levels of genetic differentiation from each other in the DAPC space, though they were still distinguishable.

**Figure 3.**
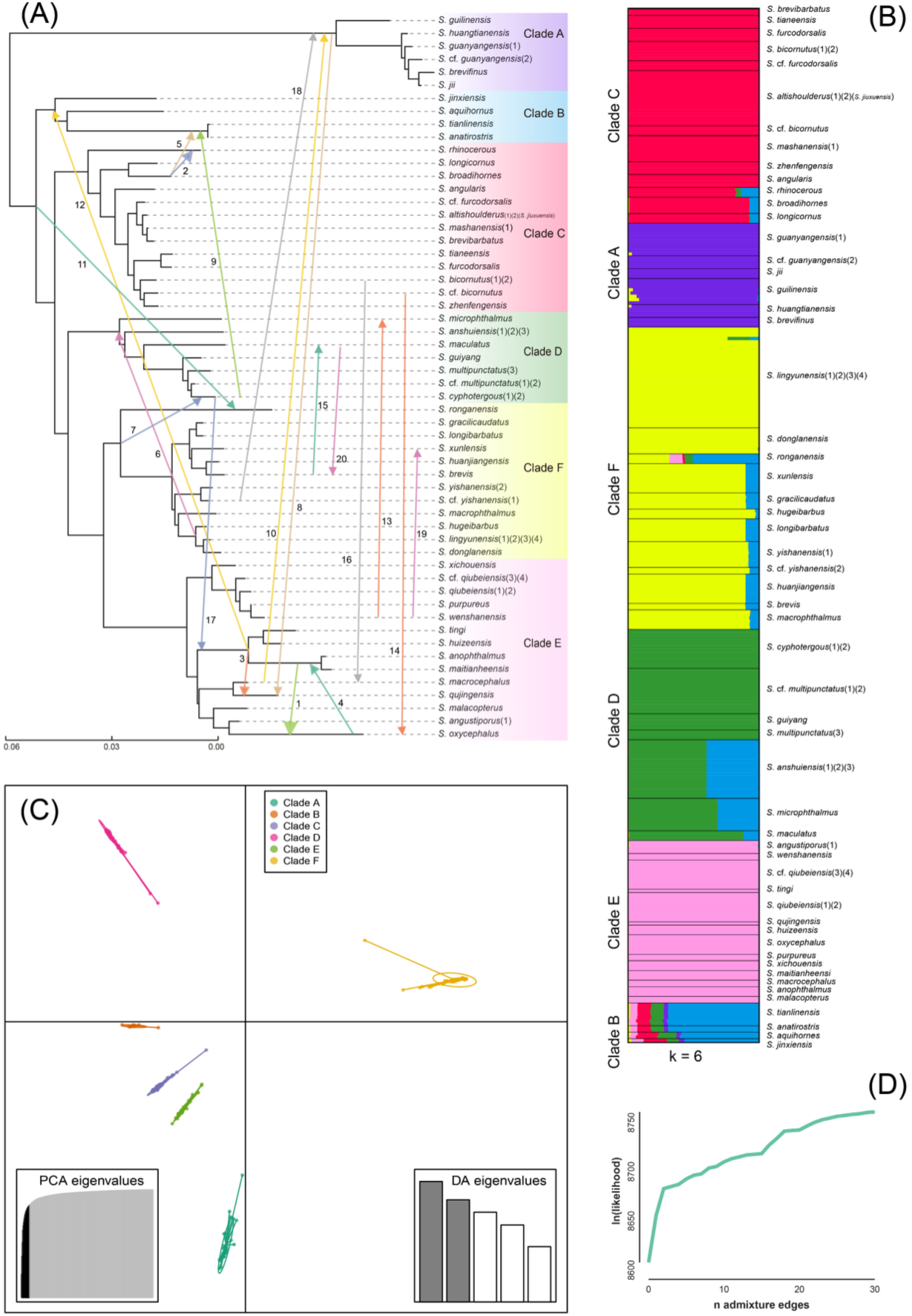
Population genetics and gene flow in *Sinocyclocheilus*. **A)** Population history of 56 *Sinocyclocheilus* species inferred with TREEMIX, including 20 migration edges. Arrows indicate the direction of gene flow, with numbers representing the intensity of migration (ranked from 1 to 20). **B)** Structure bar plot illustrating individual assignment to genetic clusters (K = 6), corresponding to the six major clades. Each bar represents one individual, and colors indicate inferred ancestry. **C)** Discriminant analysis of principal components (DAPC) plot based on SNP data showing differentiation among the six clades. **D**) TreeMix model-fit plot indicating the most likely number of admixture events based on likelihood relative increase.

#### Gene Flow and Admixture

TreeMix analyses, allowing for up to 20 migration events (m=20, where likelihood scores began to plateau; Fig. 3D), suggested several episodes of ancestral gene flow. Notable inferred events included the admixture between the ancestral lineages of (*S. anophthalmus* + *S. maitianheensis*) and *S. oxycephalus*; between *S. rhinocerous* and *S. broadihornes*; between the ancestor of (*S. anophthalmus*, *S. maitianheensis*, *S. tingi*, *S. huizeensis*) and *S. qujingensis*; and between ancestral *S. tianlinensis* + *S. anatirostris* and *S. broadihornes* (Fig. 3A). These complex admixture events may contribute to some of the topological uncertainties observed in phylogenetic reconstructions and underscore a reticulate evolutionary history in the diversification of *Sinocyclocheilus*.

### 3.4 Biogeographic Reconstructions

Among the biogeographic models evaluated, the DIVALIKE+J was the best-fitting model with the highest likelihood and the lowest Akaike Information Criterion (AICc) (Table 1). This result underscores the prominent role of founder-event speciation (i.e. jump dispersal) in shaping the current biogeographic pattern observed across the extant *Sinocyclocheilus* species. Under the DIVALIKE+J model, the genus’ range evolution involved approximately 21 dispersal events, 21 vicariance events, and five extinction events. Most of these events, particularly those associated with speciation and dispersal, occurred during the Pleistocene (Fig. 4). The ancestral range reconstruction suggests that *Sinocyclocheilus* originated in the combined Guijiang/Hejiang and Nanpanjiang River regions, likely representing the genus’ centre of origin, during the middle Miocene, around 17.96 million years ago (Ma) (Fig. 4).

**Figure 4.**
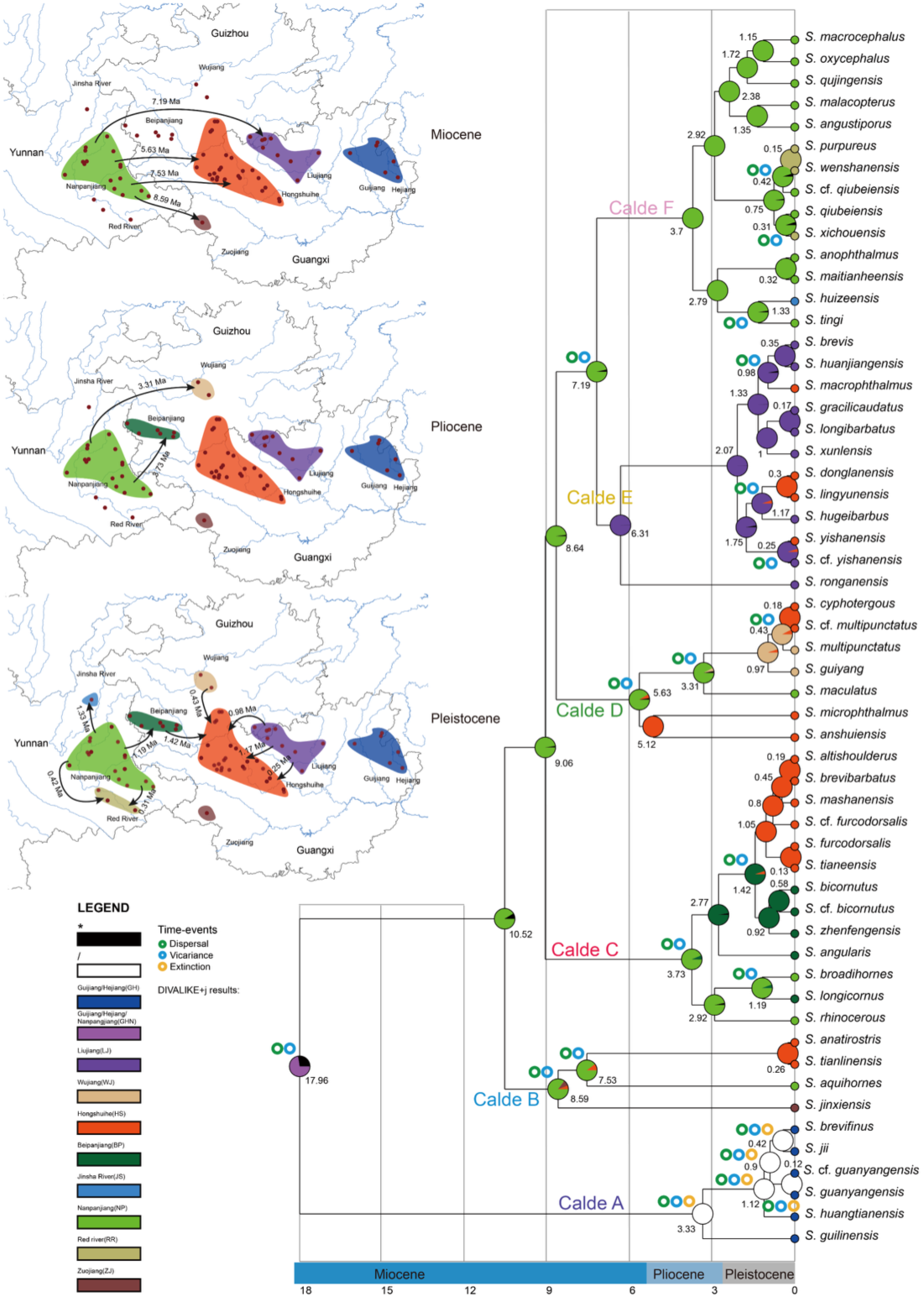
Ancestral area reconstruction of *Sinocyclocheilus*. The graphical summary (top left) illustrates changes in the genus’s distribution across the Miocene, Pliocene, and Pleistocene. Red dots mark species collection sites. Ancestral areas were inferred using the best-fit biogeographic model (DIVALIKE+J). Nine biogeographic regions were defined based on major river basins: GH: Guijiang and Hejiang; LJ: Liujiang; HS: Hongshuihe; NP: Nanpanjiang; BP: Beipanjiang; RR: Red River; JS: Jinsha River; WJ: Wujiang; ZJ: Zuojiang. Arrows indicate inferred dispersal events, with adjacent numbers denoting their estimated timing (in millions of years, Ma). Current species ranges are denoted as coloured circles at each tip of the tree, corresponding to their respective region. Pie charts at each node represent the relative probability of ancestral area assignments, with the median node ages labeled.

**Table 1.**
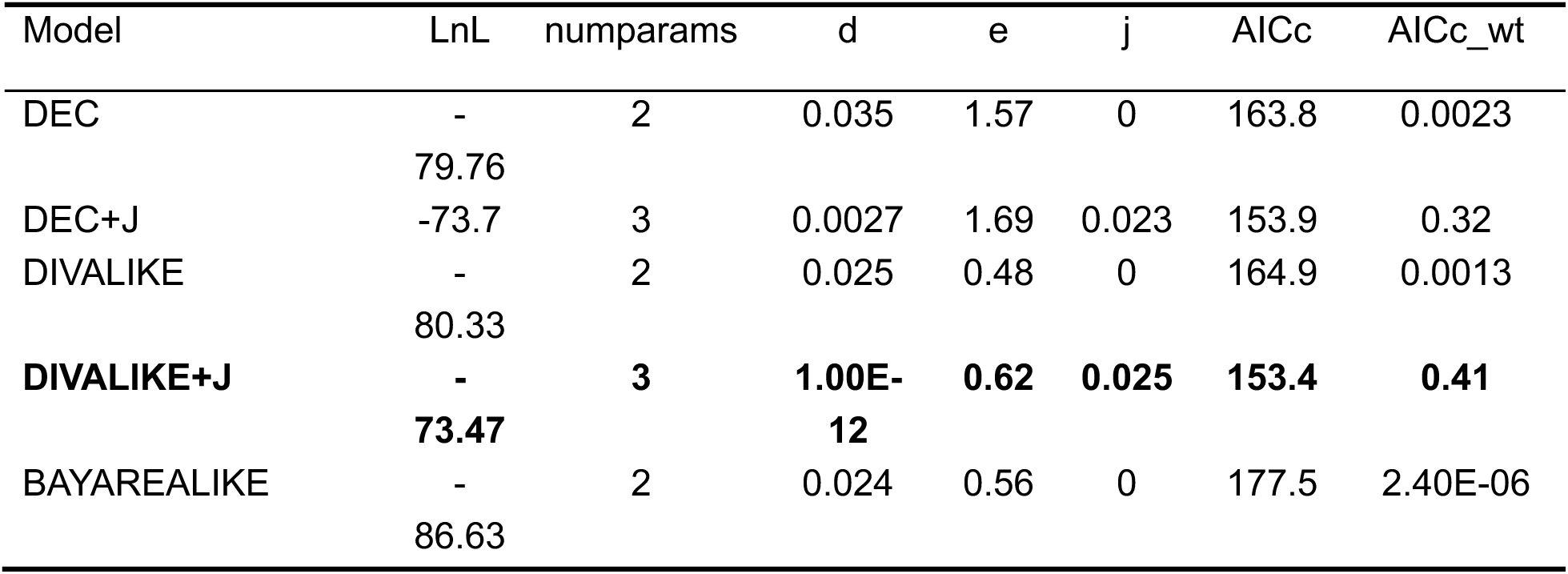
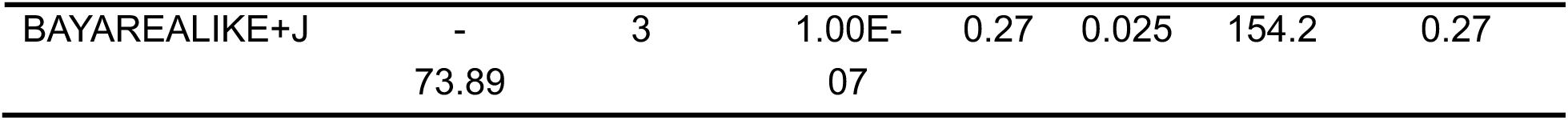
Biogeographical model comparison with and without jump dispersal events (+J) for *Sinocyclocheilus* species. Abbreviations as follows: LnL = log-likelihood; numparams = number of parameters in each model; d = dispersal rate; e = extinction rate; j = founder-event speciation rate; LRT = likelihood-ratio test; AICc = Akaike Information Criterion corrected; AICc_wt = AICc weight.

Following the initial diversification of *Sinocyclocheilus*, several key colonisation events originated from ancestral lineages in the Nanpanjiang River region. The Hongshuihe River was colonised twice, at approximately 7.53 Ma and 5.63 Ma. Other early dispersal events during the late Miocene included colonization of the Zuojiang River around 8.59 Ma and the Liujiang River around 7.19 Ma. Subsequent expansions from the Nanpanjiang River region extended into the Beipanjiang River system and into the Wujiang River system during two distinct periods in the Pliocene, around 3.73 Ma and 3.31 Ma (Fig. 4).

Subsequent dispersal events during the Pleistocene accounted for most range evolution events, likely contributing to the genus diversification (Fig. 4). For instance, in Clade C, dispersal was inferred from the Beipanjiang River to the Hongshuihe River at ∼1.42 Ma, and from the Nanpanjiang River to the Beipanjiang River at ∼1.19 Ma. In Clade D, lineages originating in the Wujiang River are inferred to have dispersed into the Hongshuihe River ∼0.43 Ma. Clade E showed three independent colonisation events from the Liujiang River to the Hongshuihe River, dated to ∼1.17 Ma, ∼0.98 Ma, and ∼0.25 Ma. In Clade F, lineages from the Nanpanjiang River expanded into the Jinsha River system around 1.33 Ma, and later colonised the Red River system during two occasions, at ∼0.42 Ma and ∼0.31 Ma. Finally, for Clade A, the model inferred a very dynamic biogeographic history shaped by five dispersal events, five vicariance events, and five extinction events.

### 3.5 Estimation of Genetic Connectivity and Barriers to Gene Flow

The FEEMS cross-validation procedure identified an optimal smoothing parameter (λ = 2.069) that minimized the L2 prediction error (Fig. 5A), and this value was used for subsequent analyses and visualization of migration surfaces. The resulting FEEMS map revealed several pronounced genetic barriers across the distribution range of *Sinocyclocheilus*, indicating regions of restricted gene flow. Strong barriers were detected in the southwestern part of Guizhou Province and the northeastern part of Yunnan Province. These regions correspond geographically to the Beipanjiang River valley and the Wumeng Mountain Range (Fig. 5B). Furthermore, low effective migration rates were also inferred in eastern Yunnan, likely reflecting the Nanpanjiang River valley acting as another geographical barrier impeding gene flow (Fig. 5B). Within the Guangxi Zhuang Autonomous Region, a distinct genetic barrier was detected in the northwestern and central areas, likely associated with limited gene flow across the Hongshuihe River valley (Fig. 5B). Migration rates were also reduced in northwestern Guangxi, potentially due to the isolating effect of the Zuojiang River valley (Fig. 5B). Similarly, the northern and northeastern regions of Guangxi exhibited very low effective migration rates, a pattern that may reflect the Guijiang and the Liujiang River valleys acting as barriers to genetic exchange (Fig. 5B).

**Figure 5.**
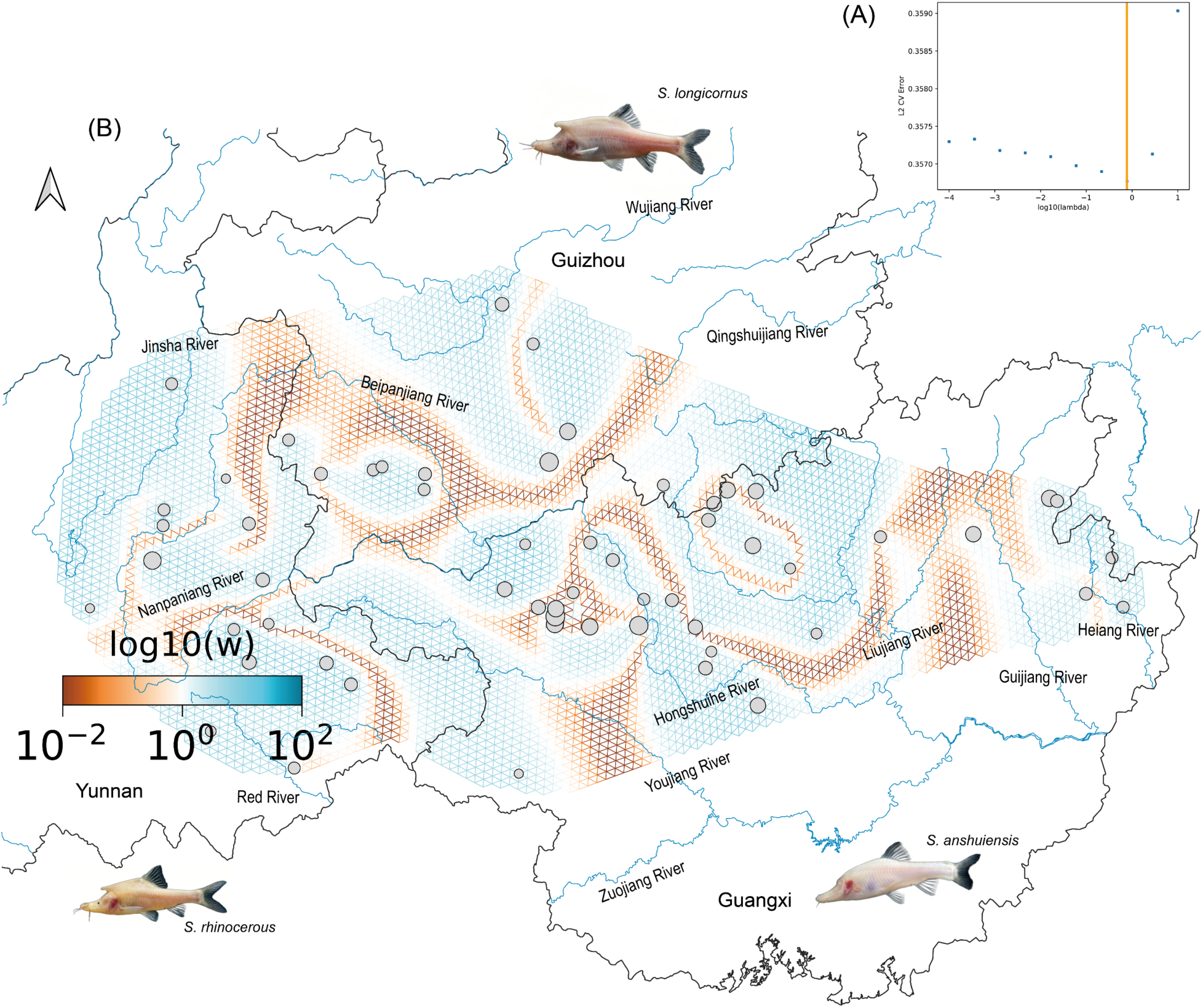
Effective migration surfaces in *Sinocyclocheilus* inferred by FEEMS. A) Cross-validation results used to select the optimal smoothing parameter (λ = 2.069) for modelling. B) Effective migration surfaces. Areas of low effective migration (genetic barriers) are depicted in brown, while regions of higher effective migration (gene flow corridors) are shown in blue. Significant genetic barriers correspond to the Beipanjiang River and Wumeng Mountains, separating populations in southwestern Guizhou and northeastern Yunnan. Additional areas of low migration rates characterizing barriers to gene flow are also evident across the Nanpanjiang, Hongshuihe, Youjiang, Guijiang, and Liujiang River valleys, highlighting their roles as impediments to contemporary gene flow.

## 4. DISCUSSION

By employing a genomic approach to examine the evolutionary history of the *Sinocyclocheilus* cavefish radiation, our study clarified its species boundaries and overcame the limitations of mitochondrial markers that previously obscured the group’s true diversity. We uncovered cryptic lineages using several complementary methods and refined the genus’s evolutionary relationships. Building on this updated species framework, we investigated both contemporary and historical processes shaping diversification. Population genomic analyses revealed patterns of genetic structure and gene flow. Divergence estimates and biogeographic reconstruction traced the origin of this radiation to the mid-Miocene, shaped by vicariance, dispersal, and environmental change. The outcome is a comprehensive evolutionary narrative of a radiation characterised by cryptic speciation, subterranean habitat specialisation, and high levels of endemism, demonstrating that *Sinocyclocheilus* diversity arose through repeated episodes of isolation and secondary contact across a dynamic Karst landscape.

### 4.1 Integrative Genomics in Resolving Complex Radiations

Adaptive radiations such as *Sinocyclocheilus* often exhibit a mismatch between morphological evolution and molecular divergence, complicated further by incomplete lineage sorting (ILS) and possible reticulate evolution, all of which hinder systematic resolution (Seehausen, 2004). Our findings conform to this pattern. While earlier studies based on morphology and limited mitochondrial markers have laid the foundation for *Sinocyclocheilus* systematics (e.g., Zhao & Zhang, 2009; Mao et al., 2021), our genome-wide dataset provides considerably improved resolution. The phylogenomic framework derived from RAD-seq data (Fig. 2) strongly supports clades that were previously weakly resolved using Sanger sequencing (Fig. 1), consistent with the view that phylogenomics can overcome the confounding effects of ILS in recent radiations (Edwards et al., 2016).

Combining genomic data with species delimitation and population genetic analyses proved highly informative. Taxa previously treated as widespread species or species complexes often resolved into multiple, genetically distinct, though sometimes morphologically cryptic, lineages under the RAD-seq + BPP framework (especially within Clades C and D). Six putative new species (*S.* cf*. guanyangensis*, *S.* cf*. multipunctatus*, *S.* cf*. bicornutus*, *S.* cf*. furcodorsalis*, *S.* cf*. qiubeiensis*, and *S.* cf*. yishanensis*) underscore this phylogenetic structure. Morphological differences are subtle and only became evident after genomic distinctiveness had been established. In the absence of genome-scale resolution, such lineages might have been dismissed as intraspecific variation. Although *Sinocyclocheilus jinxiensis* was previously assigned to *Pseudosinocyclocheilus* based on phenetic characteristics (Zhang & Zhao, 2016), our phylogeny supports its placement in *Sinocyclocheilus*(Figs. 1-2, Supplementary Figs. S2-S5 and S8-S11).

### 4.2 Methodological Congruence and Conflict

Species delimitation still remains contentious in systematics (Sites & Marshall, 2003; Carstens et al., 2013; Chambers et al., 2025; Emerson, 2025). We directly compared multiple methods applied to both Sanger and RAD-seq datasets. Sanger-based approaches (ABGD, bPTP, mPTP) consistently delimited fewer species than RAD-seq based BPP analyses, which supported the recognition of six cryptic species (*S.* cf*. guanyangensis*, *S.* cf*. multipunctatus*, *S.* cf*. bicornutus*, *S.* cf*. furcodorsalis*, *S.* cf*. qiubeiensis*, and *S.* cf*. yishanensis*). This partly reflects differences in data resolution and the limitations of single-locus and distance-based methods in resolving shallow divergences or accommodating high intraspecific variation (Papadopoulou et al., 2008; Sukumaran & Knowles, 2017).

BPP’s strength lies in its probabilistic treatment of gene tree–species tree discordance while accounting for ancestral polymorphism (Yang & Rannala, 2010). However, its results can be sensitive to priors and guide tree topology (Leaché & Fujita, 2010), and its assumption of post-divergence isolation is frequently violated in nature. Moreover, our TreeMix analysis (Fig. 3A) revealed extensive gene flow among lineages, further complicating species boundaries. We argue that concordance across MSC models, genetic structure, and phenotypic evidence provides the strongest support for species delimitation, while discrepancies should be viewed as a point of further inquiry into aspects such as the natural history of the system.

### 4.3 Geography of Speciation: Vicariance, Dispersal, and Climate

*Sinocyclocheilus* diversification reflects a complex interplay of geological isolation, episodic connectivity, and climatic shifts. Our population structure analyses (Fig. 3B, C) reveal strong genetic clustering congruent with the geographically structured phylogenomic clades, indicating that most species are genetically isolated by the region’s fragmented Karst landscape (Niemiller et al., 2013).

Biogeographic reconstruction places the origin of the radiation in the Nanpanjiang and Guijiang/Hejiang river systems during the mid-Miocene (∼18 Ma; Fig. 4), coinciding with major drainage reorganisation along the southeastern Tibetan Plateau (Ma et al., 2019; Sun et al., 2022). This initial vicariance appears to have triggered initial diversification. However, vicariance alone cannot explain the full pattern. The superior fit of the DIVALIKE+J model indicates that founder-event speciation has been recurrent. After initial separation, repeated dispersal into new river basins, facilitated by tectonic activity and climate-driven hydrological shifts, has shaped the distribution and evolution of lineages (Wan et al., 2007; Zhang, 2012). Major expansions into the Liujiang, Hongshuihe, and Wujiang basins appear linked to the late Miocene and Pliocene tectonic events (Yuan & Chen, 1993; Zhang et al., 2023).

Gene flow adds further complexity to this history. TreeMix analyses (Fig. 3A) reveal multiple admixture events, suggesting that intermittent hydrological connections enabled contact between diverging lineages. Such reticulation likely contributes to the inconsistent mapping of traits like eye reduction across the species tree. Contemporary gene flow remains constrained by geography: Our FEEMS analysis (Fig. 5B) highlights current barriers such as the Nanpanjiang and Hongshuihe valleys, reinforcing their long-term role in structuring genetic diversity.

### 4.4 Dynamics of Rapid Subterranean Diversification

The *Sinocyclocheilus* radiation, restricted to a topographically and ecologically complex region, offers a valuable model for studying rapid diversification. Cave systems promote isolation and strong divergent selection, both conducive to speciation (Porter & Crandall, 2003). The genus spans ecological extremes, from surface-dwellers with full vision to obligate cave forms lacking eyes and relying on enhanced mechanosensory traits (Zhao & Zhang, 2009). This combination of ecological specialisation and fragmented terrain likely creates a shifting mosaic of selection pressures and dispersal routes, promoting rapid diversification.

Our findings also provide a foundation for exploring the evolution and genetic basis of stygomorphic traits. The gene flow detected among cave lineages implies that speciation has not always occurred in complete isolation. Instead, parapatric or allo-parapatric modes, entailing divergence with gene flow, likely characterise much of this radiation (Sobel et al., 2010).

### 4.5 Conservation Implications in a Cryptic Radiation

Our findings have clear conservation relevance. The presence of numerous cryptic and locally endemic lineages suggests that *Sinocyclocheilus* diversity has been underestimated. Many lineages are restricted to single caves or micro-drainages, making them highly vulnerable to habitat degradation, groundwater extraction, and pollution (Luo et al., 2023).

Accurate taxonomy is important for conservation. By resolving taxonomic uncertainties and identifying new candidate species, our study provides essential data for defining Evolutionarily Significant Units (ESUs). Regions identified as centres of origin and diversification, particularly the Nanpanjiang basin, should be prioritised for conservation. And present-day genetic barriers revealed by our analyses point to isolated populations that may require separate management. Our integrated framework offers a strong basis for conservation planning, accounting for the improved taxonomic diversity of this threatened and unique cave fauna.

## 5. CONCLUSION

This study advances our understanding of the complex evolutionary history of the *Sinocyclocheilus* radiation by integrating phylogenomics, coalescent-based species delimitation, and biogeographic analysis. Our genome-wide dataset yielded a robust and highly resolved phylogeny, revealing far greater species-level diversity than previously recognised. Several cryptic lineages were identified, demonstrating that substantial genetic divergence often exists beneath subtle or convergent morphology. These findings call for a taxonomic revision and highlight the limitations of traditional markers in rapidly diversifying clades. The diversification of Sinocyclocheilus reflects a dynamic interplay of historical and contemporary processes. Our analyses suggest a mid-Miocene origin in Yunnan–Guizhou River systems, with a key vicariance event, likely linked to drainage reorganisation, acting as the initial trigger. Founder-event dispersal into emergent Karst habitats subsequently became a dominant mode of speciation. Yet, isolation was not complete; clear signals of gene flow between lineages suggest that secondary contact and introgression have also shaped the genomic landscape of the radiation.

## AUTHOR CONTRIBUTIONS

YWL, MM, TRM, and JY conceptualised the research and designed the methodology; YWL, JJZ, SPZ, TRM, JY, and MM conducted fieldwork and curated the data; YWL and TRM carried out formal analysis; YWL, TRM, and MM wrote the original draft; YWL made figures and tables; MM, MMV, MRP, and JY supervised; All authors verified. All authors reviewed and edited the draft. All authors read and approved the final manuscript.

## ACKNOWLEDGEMENTS

We thank the EED lab members for fieldwork; Cheng-Hai Fu for field work; MMV was supported as a postdoctoral fellow by Fundação de Amparo à Pesquisa do Estado de São Paulo (FAPESP #2019/08308-0 and #2023/16814-9) while working on this manuscript.

## DATA AVAILABILITY STATEMENT

Dara will be provided at a later stage after the paper is accepted for publication.

## ETHICS STATEMENT

The animal study was reviewed and approved by the Institutional Animal Care and Use Committee of Guangxi University (GXU), Nanning-China (#GXU-2024-282). The Guangxi Province Government approved field sampling; fish were sampled using trap nets, given the rarity of species, tissues from the fish were obtained mainly using fin clips without destructive sampling except for when whole specimens were required for further taxonomy work.

